# Cells and gene expression programs in the adult human heart

**DOI:** 10.1101/2020.04.03.024075

**Authors:** Monika Litviňuková, Carlos Talavera-López, Henrike Maatz, Daniel Reichart, Catherine L. Worth, Eric L. Lindberg, Masatoshi Kanda, Krzysztof Polanski, Eirini S. Fasouli, Sara Samari, Kenny Roberts, Liz Tuck, Matthias Heinig, Daniel M. DeLaughter, Barbara McDonough, Hiroko Wakimoto, Joshua M. Gorham, Emily R. Nadelmann, Krishnaa T. Mahbubani, Kourosh Saeb-Parsy, Giannino Patone, Joseph J. Boyle, Hongbo Zhang, Hao Zhang, Anissa Viveiros, Gavin Y. Oudit, Omer Bayraktar, J. G. Seidman, Christine Seidman, Michela Noseda, Norbert Hübner, Sarah A. Teichmann

## Abstract

Cardiovascular disease is the leading cause of death worldwide. Advanced insights into disease mechanisms and strategies to improve therapeutic opportunities require deeper understanding of the molecular processes of the normal heart. Knowledge of the full repertoire of cardiac cells and their gene expression profiles is a fundamental first step in this endeavor. Here, using large-scale single cell and nuclei transcriptomic profiling together with state-of-the-art analytical techniques, we characterise the adult human heart cellular landscape covering six anatomical cardiac regions (left and right atria and ventricles, apex and interventricular septum). Our results highlight the cellular heterogeneity of cardiomyocytes, pericytes and fibroblasts, revealing distinct subsets in the atria and ventricles indicative of diverse developmental origins and specialized properties. Further we define the complexity of the cardiac vascular network which includes clusters of arterial, capillary, venous, lymphatic endothelial cells and an atrial-enriched population. By comparing cardiac cells to skeletal muscle and kidney, we identify cardiac tissue resident macrophage subsets with transcriptional signatures indicative of both inflammatory and reparative phenotypes. Further, inference of cell-cell interactions highlight a macrophage-fibroblast-cardiomyocyte network that differs between atria and ventricles, and compared to skeletal muscle. We expect this reference human cardiac cell atlas to advance mechanistic studies of heart homeostasis and disease.

## Introduction

The heart is a complex organ, composed of four morphologically and functionally distinct chambers (**Figure 1A**), that perpetually pumps blood throughout our lives. Deoxygenated blood enters the right atrium and is propelled into the low pressure vascular beds of the lungs by the right ventricle. Oxygenated pulmonary blood enters the left atrium and then to the left ventricle, which propels blood across the body at systemic vascular pressures. Right and left chambers are separated by the atrial and interventricular septa and unidirectional flow is established by the atrio-ventricular valves (tricuspid and mitral) and ventricular-arterial valves (pulmonary and aortic). The heart contains an intrinsic electrophysiologic system, composed of the sinoatrial node in the right atria where depolarization begins and spreads to the atrioventricular node located at the top of the interventricular septum. This electrical impulse is then rapidly propagated by Purkinje fibers to the apex where contraction begins. Orchestration of the anatomical and functional complexity of the heart requires highly organized and heterogeneous cell populations that enable continuous contraction and relaxation across different pressures, strains, and biophysical stimuli in each chamber. The importance of these variables is reflected in the differences in wall thickness and mass (left, 116±20g; right 84±14g) of adult ventricular chambers ^1^.

**Figure 1:**
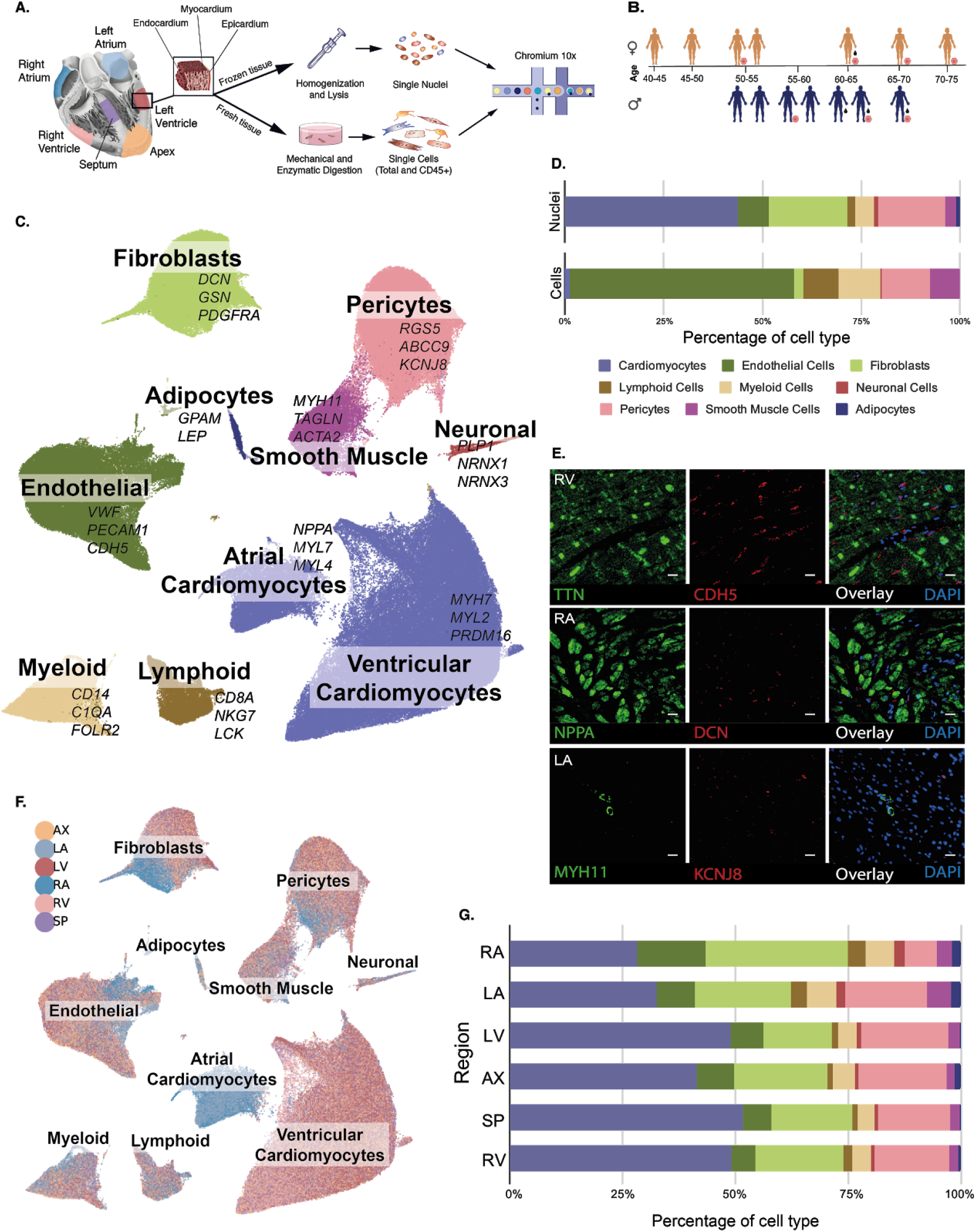
Global overview of the adult human heart cell composition. **A.** Locations of six cardiac regions sampled, including right atrium (RA), left atrium (LA), right ventricle (RV), left ventricle (LV), left ventricular apex (AX) and interventricular septum (SP). Tissues from 14 donor hearts provided 4 - 6 sites for isolation of nuclei and 7 donor hearts provided 4 - 6 tissue sites for isolation of cells. **B.** Representation of heart donors (women, top; men, bottom) denoting age, circulatory perfusion before tissue isolation (black drop sign), and donors providing both cells and nuclei (orange circle) **C.** UMAP depiction of transcriptional data from cells and nuclei that identified ten cardiac cell types cardiac tissues studied. **D.** The distribution of cell populations obtained from single nuclei (top) and cells (bottom) analyses. CD45+ cells were included in cell distributions. Colours correspond to the key in pane C. **E.** Spatial visualization of transcripts demarcating particular cell populations by smFISH with RNAscope probes was performed using fast frozen, paraffin-embedded tissue sections stained for *TTN* (green) and *CDH5* (red) in RV (top), *NPPA* (green) and *DCN* (red) in RA (middle) and *MYH11* (green) and *KCNJ8* (red) in LA (bottom). Scale bar 20μm. **F.** UMAP depiction of regional cardiac differences, parallelling panel C, and colored according to the region sampled. **G.** The distribution of cell populations, identified from all isolated nuclei within sampled regions. Colours correspond to the key in panel C.

Specialized properties of cells that enable adaptation to different biophysical stimuli in each chamber are established early in development. The heart is derived from multipotent progenitor cells residing within two heart fields. Cells of the first heart field primarily populate the left ventricle and second heart field-derived cells populate the right ventricle; both heart fields contribute to atrial cells. The distinct gene regulatory networks operant in these heart fields likely establish and prime the patterns of gene expression observed in adult cardiac cells which are further impacted by the establishment of postnatal circulation ^2^.

The cellular composition of the adult human heart, their anatomical specificities, molecular signatures, intercellular networks and spatial relationships between the various cardiac cells remain largely unknown. Single-cell and single-nuclei transcriptomics (scRNA-Seq, snRNA-seq) and multiplex smFISH imaging now enable us to address these issues at unprecedented resolution^3^. These technologies illuminate the coordinated communication of cells within their microenvironments that in the heart enable electromechanical connectivity, biophysical interactions, and autocrine/paracrine signaling required for tissue homeostasis but are perturbed in disease.

While previous studies using a combination of conventional bulk genomics and microscopy have hinted at the cellular complexity of the myocardium, limitations of these techniques have allowed definition of very few distinct cell populations ^4^. Bulk RNA-seq analysis is unable to assign gene expression to defined cell subpopulations, light microscopy fails to define features beyond morphology of cell subpopulations and immunostainings are limited to the analysis of few markers at once. Moreover, the large size of cardiomyocytes (length/width: ~100/25μM) limits the unbiased capture of single cells requiring analyses of single nuclei transcriptomics to ensure a comprehensive approach. Here, we present a broad transcriptomic census of multiple regions of the adult human heart. We profiled RNA expression of both single cells and nuclei, capturing them from six distinct cardiac anatomical regions. We also analysed the spatial distribution of selected cell populations using multiplex smFISH imaging with RNAscope probes. Our anatomically defined resolved adult human heart cell census provides a reference framework for studies directed towards understanding the cellular and molecular drivers that enable functional plasticity in response to varying physiological conditions in the normal heart, and will inform the heart’s responses to disease.

## Results

### Overview of the cellular landscape of the adult human heart

Samples were obtained from six cardiac regions including the free wall of each chamber (left/right ventricle, left/right atrium), denoted as LV, RV, LA, RA, and from the LV apex (AX) and interventricular septum (SP). To capture the heterogeneity of cardiac cell populations, samples were collected as transmural tissue segments that span the three cardiac layers (epicardium, myocardium and endocardium; **Figure 1A**) from 14 normal hearts (seven females, seven males) of North American and British organ donors (ages 40-75 years; **Figure 1B** and **Supplementary Table A1**). We isolated single cells and single nuclei, as the large sizes of cardiomyocyte (CM) are not captured by the 10X Genomics Chromium platform. Fresh tissues were mechanically and enzymatically processed to dissociate single cells, and subsequently cardiac immune cells were enriched from the cell fraction using CD45+ magnetic selection. Single nuclei were isolated from frozen heart tissues and purified by fluorescent activated cell sorting. The transcriptome of single cells and nuclei were profiled using the 10X Genomics Single Cell Gene Expression Solution (**Figure 1A**). After processing, all data from nuclei, cells and CD45+ enriched cells were batch-aligned using a generative deep variational autoencoder ^5^, prior to unsupervised clustering.

After processing and quality controls, we obtained a total of 45,870 cells, 78,023 CD45+ enriched cells and 363,213 nuclei. Overall clustering of nuclei and cells revealed ten major coarse-grained cell types based on canonical markers: atrial cardiomyocytes (aCM), ventricular cardiomyocytes (vCM), fibroblasts (FB), endothelial cells (EC), pericytes (PC), smooth muscle cells (SMC), immune (myeloid, Myel; lymphoid, Lym), adipocytes (Adi) and neural cells (NC) (**Figure 1C,E**).

Distributions of these major cell types across the six regions were estimated based on the data for nuclei only (**Figure 1D, Supplementary Table A2**). Cell proportions within ventricular regions are similar to each other (AX, SP, LV, RV), and vCM are the largest population, constituting 47.1%, followed by FB (19.5%), mural cells (PC and SMC, 18.7%) and EC (7.2%) (**Figure 1G**). The same trend was seen in both atria. The proportions of CM, EC, SMC, NC and Adi were significantly different in atrial and ventricular tissues, with CM being the largest population again, comprising 31.1% of atrial and 47.1% of ventricular cells (*p-value* for ventricular enrichment <1e-8).

We found differences in the distribution of cell types across donors, even within the same region, and correlations between different cell types at the same site (**Supplementary Table A3**). For example, in LV, AX, SP and RV tissues, the proportions of vCM and FB were negatively correlated (*p*-value=2.0e-4), while there was a positive correlation among the proportions of PC, SMC, and NC cells (*p*-value<1e-4). We suggest that these data reflect random sampling that included vessels with EC, PC, SMC, and concurrently fewer CM. However, the observed correlation between NC, SMC and PC implies a potential functional organization.

The cell distributions were generally similar in tissues of male and female hearts, but left ventricular regions (AX, SP, LV) from female donors had significantly higher mean percentages of CM (46.7±12%) compared to male donors (33.9±10%; *p*-value=0.008). This is unexpected given the average smaller heart mass of women and may reflect the small donor pool. If replicated, these data may explain higher cardiac stroke volume in women ^6^ and lower rates of cardiovascular disease.

### Inter- and intra-cardiac chamber heterogeneity of cardiomyocytes

CM are responsible for generating the contractile force of the heart, and are the most abundant cell type in the LA, RA, RV and all LV regions captured by single nuclei profiling. Transcriptional analyses of CM nuclei were identified based on high-level expression of genes encoding proteins in the sarcomere (*TTN*, *MYBPC3, TNNT2*), and involved in calcium-mediated excitation-contraction processes (*RYR2, PLN*, *SLC8A1*). Consistent with their developmental origins, functional differences in electrophysiological, contractile, and secretory processes described in previous tissue RNAseq data analyses ^7,8^, we observed distinct transcriptional signatures in atrial (aCM) and ventricular (vCM) nuclei (**Figure 1C**).

Of 10,228 genes expressed among all CM, 389 genes showed >3-fold higher expression in aCM (**Supplementary Table B1**), including *HCN1*, *MYH6*, *MYL4* and *NPPA*. In addition, we identified a higher aCM expression of *ALDH1A2* (9-fold increased), the catalytic enzyme required for synthesis of retinoic acid, that may reflect aCM derived from the second heart field ^9^. aCM also showed higher levels of *ROR2* (22-fold increased), which participates in cardiomyocyte differentiation via Wnt-signaling, as well as *SYNPR* (29-fold increased), a synaptic vesicle membrane protein with functions in TRP-channel mechanosensing by atrial volume receptors ^10,11^.

In contrast, vCM expressed 79 genes with 3-fold higher expression than aCM (**Supplementary Table B1**). Genes with significantly enriched expression included prototypic sarcomere protein genes *MYH7*, *MYL2*, and transcription factors (*IRX3, IRX5, IRX6, MASP1, HEY2*). vCM also had 20-fold higher expression of *PRDM16* than aCM, which harbors damaging variants associated with LV non-compaction ^12^. Similarly highly expressed were *PCDH7* (**Supplementary Figure B1A,B**), a molecule with strong calcium-dependent adhesive properties and *SMYD2*, which promotes the formation of protein stabilizing complexes in the Z-disc and I-band of sarcomeres ^13,14^. Expression of these genes is likely to promote tissue integrity under the conditions of high ventricular pressure and strain.

Clustering of vCM data identified five subpopulations (**Figure 2A**), with variable proportions in LV, AX, SP and RV, indicative of inter-chamber heterogeneity. vCM1 and vCM2 constituted the majority of CM (~70-80%). Subpopulation vCM1 comprised ~60% of vCM across all left ventricular sites, but only 33% of RV (**Figure 2C, Supplementary Table B2**). Small numbers of these nuclei were also detected in atrial tissues (**Figure 2B,D**). In contrast, vCM2 cells were enriched in RV (42%) compared to 11% in LV. We noted only small differences (<3-fold) in the low level of expression of some genes found in vCM1 and vCM2 (**Supplementary Table B1**), which indicates shared gene programs between the left (enriched in vCM1) and right (enriched in vCM2) ventricles.

*PRELID2* (**Figure 2E, Supplementary Figure B1B**) was ~2-fold higher in RV, as previously reported, and highest in vCM2 (**Supplementary Table B3**), supporting a biological basis for this subpopulation ^8^. Although cardiac functions of *PRELID2* remain unknown, it is developmentally expressed during mid-gestation ^15^. Using smFISH of four markers simultaneously in tissue sections, we confirmed enriched *PRELID2* expression in RV (**Figure 2G**). In comparison to other subpopulations, vCM2 had highest expression of *MYH6* and *CDH13*, a cell surface T-cadherin receptor for the cardioprotective hormone adiponectin and low density lipoproteins, both of which are associated with multiple cardiometabolic traits ^16,17^.

We also identified two subpopulations (vCM3 and vCM4) across all ventricular regions. The transcriptional profile of vCM3 was remarkably similar to a prominent RA subpopulation (aCM3, discussed below), and suggestive that these are derived from the second heart field^18^. These cells had higher levels of transcripts associated with retinoic-acid responsive smooth muscle cell genes, including *MYH9*, *CNN1* ^19,20^, and *NEXN*. vCM3 also expressed stress-response genes including *ANKRD1^21^*, *FHL1* ^22^, *DUSP27^23^*, *XIRP1 and XIRP2*. The XIRP proteins interact with cardiac ion channel proteins Nav1.5 and Kv1.5 within intercalated discs, and have been implicated in lethal cardiac arrhythmias prevalent in cardiomyopathies ^26,24^.

vCM4 contributed 6-10% to vCM populations, and expressed nuclear-encoded mitochondrial genes (*NDUFB11*, *NDUFA4*, *COX7C*, and *COX5B*; **Figure 2E**) suggestive of a high energetic state. Indeed, Gene Ontology analyses of vCM4 transcripts identified significant terms of “ATP metabolic process” and “oxidative phosphorylation” (**Supplementary Figure B1C**). These CM also had high levels of *CRYAB* encoding a heat shock protein with cytoprotective roles and antioxidant responses by CM ^25^. With concomitant high expression of genes encoding sarcomere components (**Figure 2E**) and *PLN* (**Supplementary Table B3)**, we deduced that these vCM are outfitted to perform higher workload than other vCM. Unlike scRNA-Seq analysis that identified prominent RV expression of *PLN* in embryonic mouse hearts ^26^, vCM4 were similar in both ventricles.

A small proportion (~1%) of cells comprised vCM5 and expressed high levels of *DLC1* and *EBF2 ^27^*. These molecules participate in regulating brown adipocyte differentiation and may be involved in cardiac pacemaker activity. In addition, vCM5 nuclei had higher levels of transcripts also expressed in neural lineages (*SOX5, EBF1*, and *KCNAB1*). Notably, mice with deleted *EBF1* have profound hypoplasia of the ventricular conduction system ^28^. Further confirmation and investigation of this subpopulation is required, given the small number of cells and shared marker genes with other cell types.

We identified six subpopulations of aCM, indicating considerable heterogeneity (**Figure 2B**) particularly between the right and left chambers. Notably, *HAMP*, a master regulator of iron homeostasis, was significantly enriched in over 50% of RA CM compared to 3% LA CM (**Supplementary Table B4**) ^29^, consistent with prior studies of RA tissues ^8^ and confirmed by smFISH (**Figure 2G**). *HAMP* has unknown roles in cardiac biology, but *Hamp-*null mice have electron transport chain deficits and lethal cardiomyopathy ^30^. The RA enrichment of *HAMP* may imply energetic differences between right and left aCM.

aCM1 cells comprised 68% of LA CM (**Figure 2D, Supplementary Table B2**). Transcripts from these cells encoded prototypic atrial proteins and also low levels of molecules with recognized neural functions (*ADGRL2, CHRM2, GABRB2, NFXL1, KCNMB1*, *ROBO2*). As aCM1 had no defining gene enrichment in comparison to other aCM subpopulations, we suggest they likely represent a basal aCM gene program.

aCM2 cells were more prominent in the RA than LA. Consistent with this, aCM2 cells had the highest proportion (65%, **Supplementary Table B3**) of *HAMP*-positive CM. Other enriched transcripts in this subpopulation included *SLIT3*, encoding the predominant developmental ligand for ROBO receptors in the heart ^31^, *ALDH1A2* ^9^ and *BRINP3*, involved in retinoic acid signaling, and *GRXCR2*, a molecule that supports cilia involved in mechanosensing ^32^.

As noted above, aCM3 shared a remarkably similar transcriptional profile with vCM3 including the smooth muscle cell gene *CNN1*, which we confirmed by smFISH with RNAscope probes (**Figure 2G, Supplementary Figure B1A**). The transcriptional profiles of aCM2, aCM3 and vCM3 likely indicate their derivation from the second heart field that forms the right heart chambers and associated vascular structures ^18^.

Small cell numbers comprised aCM4-6. aCM4 cells contained several transcripts (*CKM, NDUFA4, COX4I1, FABP3, HSBP1*) denoting high metabolic activity, similar to vCM4 (**Figure 2E, F**) and had the highest *NPPA* expression. aCM5 nuclei contained ventricular transcripts, including *MYH7* and *MYL2* (**Figure 2F**), confirming a previous observation ^33^. These cells had enriched *PPFIBP1* expression which participates in cell adhesion and migration ^34^ and cardioprotective *FHL2* (~5-fold higher than other aCM). aCM6 expressed transcripts (*DLC1*, *PLA2G5*, *GSN*) associated with stromal cells similar to vCM5.

The recent identification of cardiac injury and decompensation in 10-20% of COVID-19 patients prompted us to analyze cardiac expression of the viral receptor, *ACE2* and proteases that prime viral entry (*TMPRSS2*, *CTSB*, *CTSL*) ^35^. *ACE2* expression was low in vCM, albeit 2-fold higher than in aCM, and highest in vCM3, that is RV enriched. CM do not express *TMPRSS2* but have high levels of *CTSB* and *CTSL* (**Supplementary Figure A3**).

**Figure 2: CM cells of the human heart. A.** UMAP depiction of transcription data identified five vCM subpopulations **B.** UMAP depiction of transcription data identified six aCM subpopulations. **C.** Representation of the distributions of vCM subpopulations (identified in A) within LV, RV, AX, and SP regions. **D.** Representation of the regional distributions of aCM subpopulations (identified in B) within LA and RA regions. **E.** Dot plot of the expression of selective marker genes in vCM subpopulations. **F.** Dot plot of the expression of selective marker genes in aCM subpopulations. **G.** Multiplex smFISH with RNAscope probes yields *in situ* visualization of transcripts enriched in CM subpopulations. **G.I.** RV expression of *TNNT2* (green) and *PRELID2* (red; marked with stars). **G.II.** LV expression of *TNNT2* (green) and *FHL1* (red). **G.III.** RA expression of *TNNT2* (green) and *HAMP* (red). **G.IV.** RA expression of *TNNT2* (green) and *CNN1* (red) i. Nuclei were DAPI-stained (dark blue). Scale bar 10μm.

### Vascular cells of the heart

The vascular cellular compartment including EC, SMC, PC,analysed alongside mesothelial cells, separated into 15 different cell populations with clear anatomical and arterio-venous specificities (**Figure 3A**). Single cell RNA-seq were particularly useful here, in addition to single nuclei, providing half of the transcriptional data (**Supplementary Table C1**) and enabling analyses of putative (trans)differentiation relationships using RNA velocity (**Supplementary Figure C1**).

**Figure 3:**
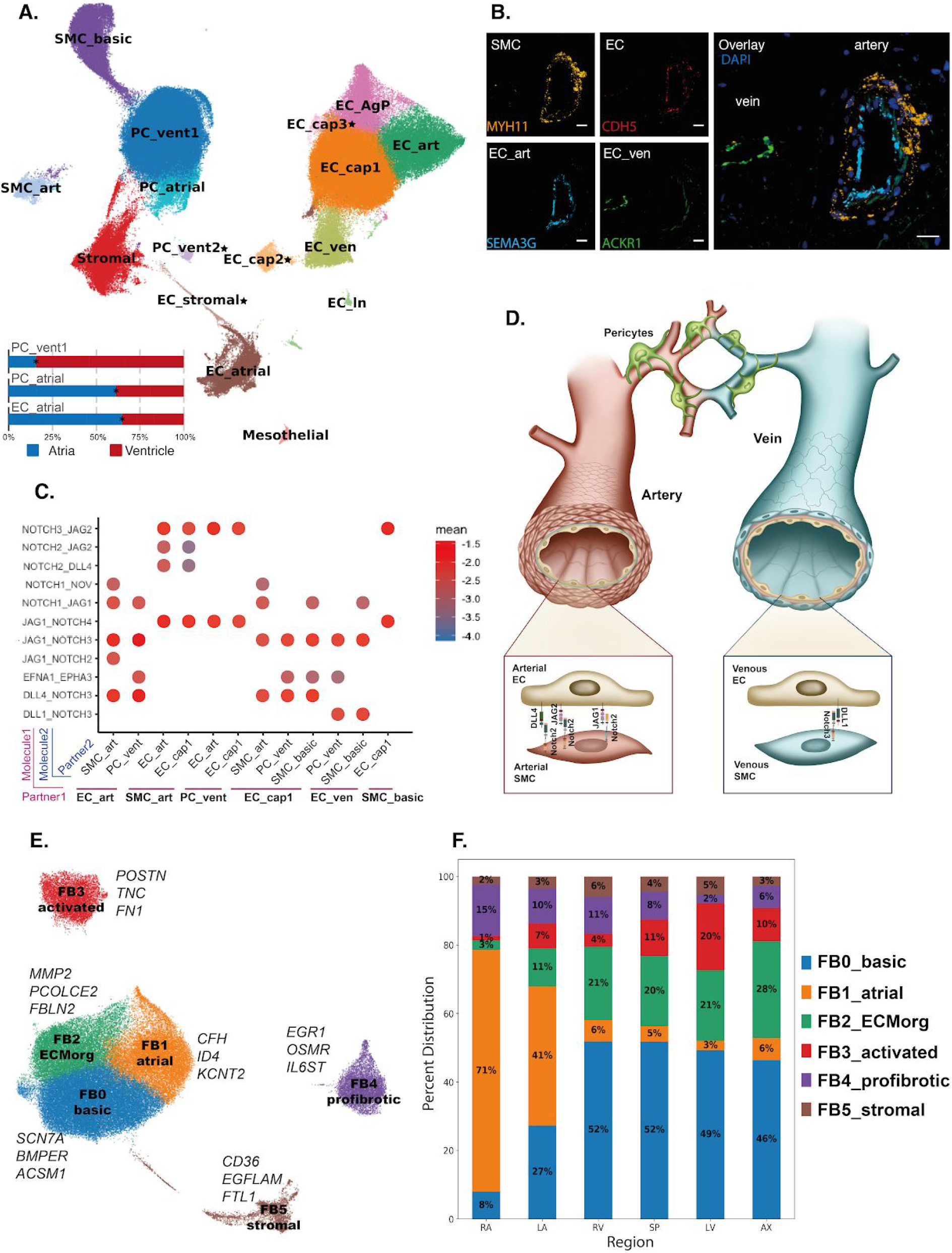
Stromal cells of the human heart. **A.** Visualization of transcriptional data from vascular and mesothelial cells by UMAP. Clustering identifies 16 subpopulations. Bar graphs represent the statistically significant distribution of the indicated PC and EC populations in the atrial (RA and LA) and ventricular (AX, LV, RV, SP) anatomical regions, using nuclei only (EC_cap = capillary endothelial cells, EC_art = arterial endothelial cells, EC_ven = venous endothelial cells, EC_atrial = atrial endothelial cells, EC_ln = lymphatic endothelial cells, PC_vent = ventricular pericytes, PC_atrial = atrial pericytes, Stromal, EC_stromal = stromal cells, SMC_basic = smooth muscle cells, SMC_art = arterial smooth muscle cells, Mesothelial = mesothelial cells), EC_cap2, EC_cap3, PC_ven2 and EC_stromal (denoted*), were represented in only a subset of donors and/or predominant contribution from nuclei or cells. **B.** Staining of myocardium from AX including an artery and vein by smFISH with RNAscope probes. Images show specific expression of *MYH11* in vascular SMC (thick in artery and very thin in the small caliber vein shown), *CDH5* in the endothelial layer, while *SEMA3G* and *ACKR1* are expressed respectively in arterial and venous EC. Nuclei were counterstained with DAPI (dark blue). Scale bar 20μm. **C.** Selected ligand-receptor interactions show specificity of NOTCH ligands-receptors pairing in defined vasculature districts and a specific putative EFNA1-EPHA3 interaction between EC-PC. Color indicates the mean expression level of interacting molecule in partner 1 and interacting molecule partner 2. (CellPhoneDB *p*-value of the specificity of the interactions=10e-5.) **D.** Illustration of the anatomical relationship of SMC, PC and EC in arteries and veins. The zooms depict the selected receptors and ligands identified in panel **C**. **E.** UMAP representation of transcriptional data from FB identified six subpopulations. **F.** Regional distributions (mean abundance) of FB subpopulations (identified in **E**) within LA, RA, LV, RV, AX and SP regions. Regional specification of fibroblasts was shown by the separation of ventricular- and atrial-enriched clusters.

Endothelial cells, defined by pan-EC markers *PECAM1*, *CDH5* and *VWF*, were highly heterogeneous, consisting of eight subpopulations (**Figure 3A**). The most abundant subpopulation (53.0% of all EC) was annotated as capillary EC (EC_cap) given the enrichment of *RGCC* and *CA4*, marker genes of microvascular endothelium in the mouse endothelial cell atlas ^36^. A separate cluster, expressing endothelial capillary markers, was largely represented by cells (**Supplementary Figure C2**), and enriched for antigen-presentation and immune-regulation related transcripts (EC_AgP) including *CXCL1/3*, *CCL2*, *IL6* and *ICAM1^37^*, as well as *FOS* and *JUN* that can be induced in a variety of situations including cellular stress. Neighbouring clusters contained an arterial endothelial subpopulation (EC_art), as confirmed by specific enrichment of *SEMA3G*, *EFNB2*, *DLL4*, and a venous endothelial subpopulation (EC_ven) expressing a well-accepted venous marker *NR2F2^38^ and ACKR1* (**Figure 3A**, **Supplementary Figure C2 and Supplementary Table C2**). *ACKR1* was reported to also be expressed in endothelial stalk cells in cancer^39^, EC expanding in cirrhosis^40^ and the endothelium of lymphnodes’ post-capillary venules^41^. Using a combination of pan-EC (*CDH5)*, arterial EC (*SEMA3G*), and smooth muscle cells (*MYH11)* markers, we found *ACKR1* expression in the EC of small caliber veins (as defined by a *CDH5+* SEMA3G-EC with thin SMC layer) (**Figure 3B**).

Of note, one EC cluster enriched for atrial-derived cells (EC_atrial) and specifically expresses at least two genes previously used as endocardial markers: *SMOC1*, a newly proposed endocardial marker ^42^ whose product regulates angiogenesis ^43^; *NPR3*, enriched in mouse endocardium as shown in fate-mapping studies ^44^. Lymphatic EC (EC_ln) were defined by expression of *PROX1*, *TBX1* and *PDPN*, representing ~1% of the EC (**Supplementary Figure C2**).

Two EC clusters expressing capillary markers were represented by a minority of donors. EC_cap3 was mostly represented by three donors and EC_cap2 is a small cluster deriving almost exclusively from one heart (**Supplementary Figure C2**), thus their significance remains unclear.

We captured a small subpopulation of mesothelial cells expressing *MSLN*, *WT1* and *BNC1*, but not EC, FB or mural lineage genes, indicating that these are likely epicardial cells ^45,46,47^. *In situ* validation showed BNC1+/CDH5− subpopulation of mesothelial cells lining the epicardium (**Supplementary Figure C4**).

Pericytes, defined by expression of *ABCC9* and *KCNJ8* showed strong regionality with ventricular-enriched (PC_ven1/2) and atrial-enriched (PC_atrial) clusters. Even though the expression profiles of ventricular and atrial PCs overlapped, PC_ven showed higher expression of *NCAM2* encoding for an adhesion molecule, *CD38* a transcript for a multifunctional receptor with ectoenzyme activity and involvement in cell adhesion, as well as *CSPG4*, a proteoglycan with an essential role in microvascular morphogenesis and cross-talk with EC ^48^. One small cluster (PC_vent2) was largely represented by just three donors and showed expression of genes indicative of oxidative stress (**Figure 3A, Supplementary Figure C3 and Supplementary Table C2**). Notably, PC expressed the highest levels of *ACE2*. *TMPRSS2* is not detected in these cells although cathepsin B and L (CTSB, CTSL) are expressed albeit at low levels (**Supplementary Figure A3**).

Smooth muscle cells showed two distinct subpopulations, arterial (SMC_art), expressing contractile *CNN1^49^* and high levels of *ACTA2*, and likely venous (SMC_basic) with more immature features expressing *LGR6*, previously described in the lung ^50^ and *RGS5*, related to SMC proliferation ^51^. (**Figure 3A, Supplementary Figure C3 and Supplementary Table C2**). This is in line with in vitro models predicting that while arterial SMC are more contractile, the venous seem more dedifferentiated and potentially proliferative ^52^.

Cell-cell interaction analyses using CellPhoneDB predicted heterotypic connections between EC_art and SMC_art (**Figure 3C, D**). We also predicted differences between the interconnectivity of arterial (EC_art and SMC_art) *versus* venous (EC_ven and SMC_basic) and smaller vessels (PC-EC_cap), including specific patterns of *Notch*-ligand interactions (**Figure 3C, D and Supplementary Table C3**)^53^.

Among vascular populations, we identified stromal (Stromal3) cells with expression of PC and EC markers, including *RGS5*, *ABCC9*, *VWF*, and *SOX18*. Using RNA velocity analyses, we observed a directionality implying that Stromal3 may represent a transitional state between PC and EC (**Supplementary Table C2, Supplementary Figure C1**). Transdifferentiation of stromal populations has been described but remains controversial in the literature ^54 55 56^.

### Cardiac fibroblast cells

We identified FB based on enriched expression of *DCN*, *GSN* and *PDGFRA*. Subclustering on FB revealed six subpopulations (**Figure 3E**) with regional enrichment in ventricle (FB0_basic; 2.8-fold higher) and atria (FB1_atrial; 6.5-fold higher). These findings are consistent with different specific functional properties of atrial FB, including stronger profibrotic responses ^57, 58^. As genes expressed in FB0_basic and FB1_atrial were found in all ventricular or atrial FB, respectively, we suggest these define a basal, chamber-specific FB gene program (**Supplementary Figure D1A, Supplementary Table D1**).

Three additional subpopulations (FB2_ECMorg, FB3_activated, FB4_profibrotic) were identified across all regions, albeit in variable proportions (**Figure 3F**). In comparison to other cardiac regions, FB2_ECMorg and FB3_activated were less abundant (0.85-fold) in the right atria, and FB4 was less abundant (2.5-fold) in all left ventricular sites (**Supplementary Figure D1C**). FB2_ECMorg showed higher expression of genes involved in extracellular matrix (ECM) production, remodeling, and cleavage including *FBLN2, PCOLCE2*, *MFAP4* and *MFAP5 (*myofibrillar-integrin linker proteins), and *MMP2* (matrix metalloproteinase 2). FB3_activated expressed genes responsive to TGF-*β* signaling (*POSTN*, *CTGF*, *TNC* and *RUNX1)*. In contrast, FB4_profibrotic had lower expression levels of ECM proteins but higher expression of cytokine receptors (*OSMR, IL6ST)*, transcription factors (*JUNB*, *EGR1*, *JUN*, and *HIF1A)* and other genes in the OSM pathway (**Supplementary Figure D1B, Supplementary Table D2**) ^59, 60^. As macrophage secretion of oncostatin M, the OSMR and IL6ST ligand, would suppress cardiac fibrosis ^61^, the distinctive gene programs of FB subpopulations are likely to govern stress-responsive cardiac remodeling. FB5_stromal expressed multiple stromal markers, including *CD36*, *EGFLAM* and *FLT1*.

To better identify subpopulations independent of region-specific basal gene expression, we analyzed atrial (aFB) and ventricular (vFB) fibroblasts separately (**Supplementary Figure D1D-G**). These analyses recapitulated the FB subpopulations described above, including FB4_profibrotic cells in each chamber that express *OSMR* and *IL6ST*, and also identified another ECM-producing FB that did not emerge from the integrated analysis of all chambers. These aFB2 and vFB2 populations differed in their expression of collagen isoforms and other ECM proteins (**Supplementary Figure D1H, Supplementary Table D3-4**). Similarly, we identified chamber-specific aFB4 and vFB3 that differed in the expression of proteins involved in organising connective tissue.

### Immune cells and cardiac repair

Immune cells are known to play key roles in both cardiac homeostasis, as well as inflammation, repair and remodeling; however, they were unrepresented in our single nuclei dataset. Therefore, we used the pan-immune marker CD45 to enrich for this cell population. We defined cardiac-resident *versus* circulating cells by calculating the enrichment in perfused tissue *versus* non-perfused tissue (i.e. NRP *versus* non-NRP donors), as well as by mapping markers associated with tissue residence.

Myeloid cells represented around 5% of the total cardiac cells and comprised eight distinct subpopulations (**Figure 4A** and **Supplementary Figure E1A**), characterised by well-defined gene markers (**Supplementary Table E1**). The largest myeloid subpopulation were M2 macrophages (MØM2_CD163L1+ = 10582 cells) followed by CD52+ monocytes (Mono_CD52+ = 5120 cells). Two of the predicted yolk sac derived (*CCR2-*), resident macrophages, MØM2_trFOLR2+ and MØ_trTMSB4X+, with similar transcriptional profiles, expressed *TMSB4X*, *FOLR2*, *S100A9*, *HLA-DRB1*, *HLA-DRA* and *LYVE1*, which partially resemble resident myeloid cells described in mice and implicated in cardiac remodelling ^62^; however, unlike the murine model, the human cells did not express *TIMD4* (**Supplementary Figure E**1A and **Supplementary Table E1**). MØM2_trFOLR2+, dendritic cells (DC_class1) and MØ_trTMSB4X+ expressed the anti-inflammatory genes *GRN*, *HIF1A*, *AIF1* and *TIMP1* ^63^ (**Supplementary Figure E1C**), suggesting that these populations could be associated with cardiac homeostasis ^64^.

**Figure 4:**
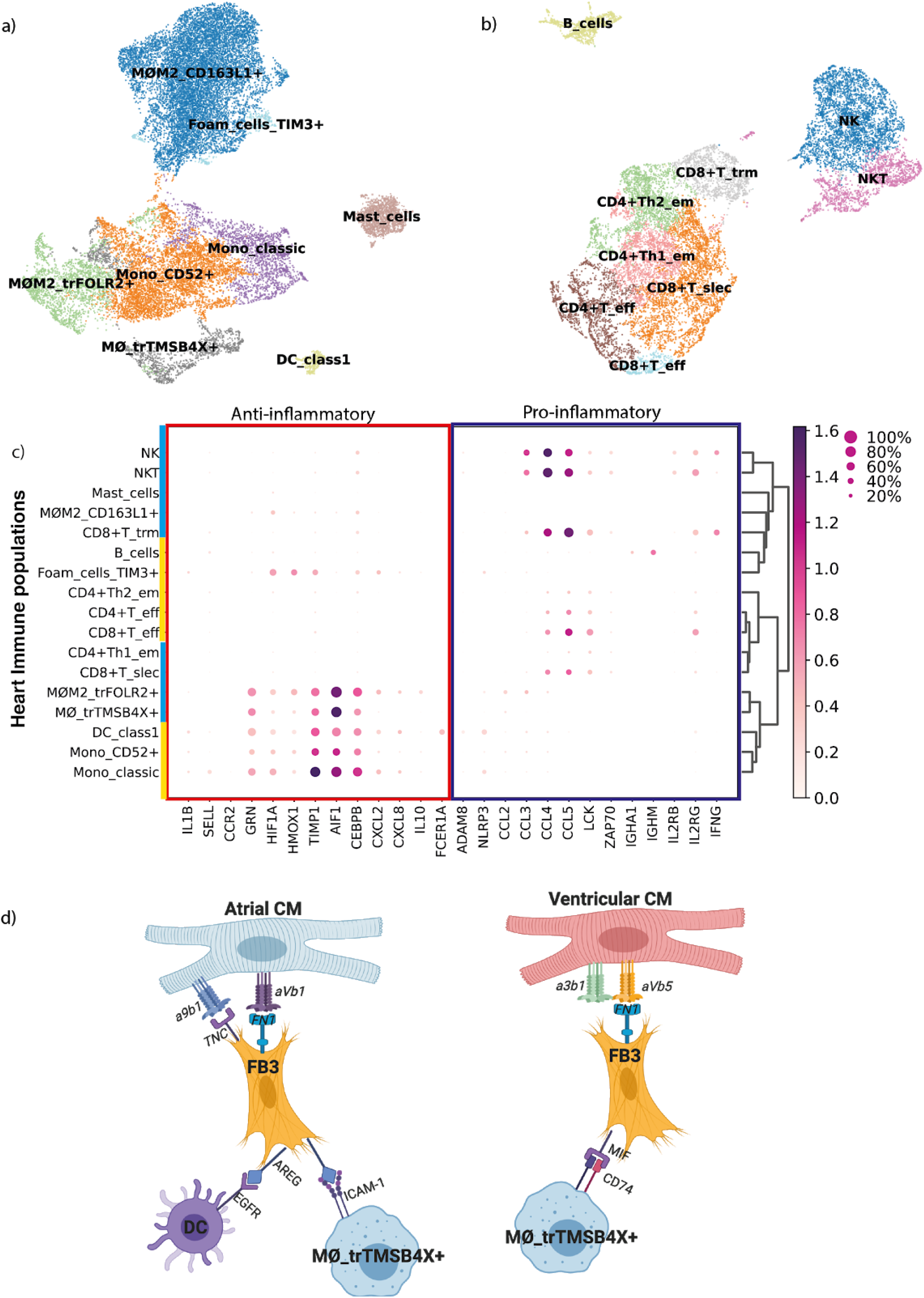
Cardiac-resident immune populations and cell-cell interactions. **A.** Cardiac resident myeloid subpopulations (listed by prevalence) included *CD163L1*+ M2 macrophages (MØM2_*CD163L1*+), *CD52*+ monocytes (Mono_*CD52*+), FOLR2+ M2 tissue-resident macrophages (MØM2_tr*FOLR2*+), Classical monocytes (Mono_classic), *TMSB4X*+ tissue-resident macrophages (MØ_tr*TMSB4X+*), mast cells, *TIM3+* foam cells (Foam_cells_*TIM3+*) and class 1 dendritic cells (DC_class1)**. B.** The most prevalent resident lymphoid cells were natural killer cells group (NK) and short-lived effector cells CD8+ T cells (CD8+T_slec) followed by effector-memory Th1 CD4+ T-cells (CD4+Th1_em), effector-memory Th2 CD4+ T-cells (CD4+Th2_em), effector CD4+ T-cells (CD4+T_eff), Natural Killer T-cells (NKT), tissue-resident memory CD8+ T-cells (CD8+T_trm), B cells (B_cells) and effector CD8+ T-cells (CD8+T_eff). **C.** Cytokines and chemokines associated with anti-(red) and pro-inflammatory (blue) processes in myeloid (golden) and lymphoid (cerulean) populations. Log fold change (logFC) in gene expression is indicated by the colour scale. Dot size represents the proportion of a cell population expressing a given gene. **D.** BioRender representation of predicted receptor-ligand interactions (based on CellPhoneDB.org) across cells with relevance to cardiac remodeling. Signalling between aCM and activated FB3 is mediated through *TNC_a9b1* and *FN1_aVb1* receptor-ligand complexes. FB3-DC interaction involves the *EGFR - AREG* receptor-ligand complex, and FB3 - MØ_tr*TMSB4X*+ via *ICAM1-AREG receptor-ligand complex*. vCM and FB3 are predicted to interact *via* the *FN1_a3b1* or *FN1_aVb5* complexes. FB3 is predicted to interact with MØ_tr*TMSB4X*+ *via* the *CD74-MIF* complex.

The cardiac-resident lymphoid compartment was composed of nine subpopulations (**Figure 4B** and **Supplementary Figure E1B**), with the largest being NK cells, followed by short-lived effector *CD8*+ T-cells (CD8+T_slec), activated CD4+ T-cells (CD4+T_eff) and effector memory Th2 CD4+ T-cells (**Supplementary Table E2)**. NK, NKT, effector CD8+ T-cells (CD8+T_eff), CD4+T_eff and CD8+T_slec transcriptomes were enriched for genes encoding pro-inflammatory chemokines (*CCL3*, *CCL4*, *CCL5), IL2RG*, and *LCK*, encoding a T-cell antigen receptor-linked signal transduction protein whose activation yields lymphokine production ^65^.

To evaluate cell - cell interactions of the immune cells in cardiac homeostasis and remodelling we used cellphonedb ^66^. Thus we predicted putative cross-talk among myeloid and lymphoid cells, CM, and FB (**Figure 4C** and **Supplementary Table E3**) ^64^. This analysis predicted that DC and MØ_trTMSB4X+ interacted with FB3 in a distinct manner in atria and ventricles, as shown in **Figure 4D**. The FB3 subpopulation signaled to aCM and vCM in turn, forming a cellular circuitry which may be relevant for healthy cardiac homeostasis.

### Neural populations

The heart is innervated by both sympathetic and parasympathetic components of the autonomic nervous system, which reside in ganglionated plexuses on the epicardial surface and contribute to regulation of heart rhythm ^67^. Neural cells (NC) innervate the sinoatrial and atrioventricular nodes, from which an activating wave-front propagates throughout myocardial tissue *via* nerve bundles and Purkinje fibers. We identified 3485 NC from cardiac tissues, predominantly (60%) from left chambers, presumably due to multiple left ventricular sites. All NC expressed transcripts (*NRXN1, NRXN3, KCNMB4*) typically found in the central nervous and cardiac conduction system including sodium and potassium channels ^68^ (**Supplementary Table F1)**.

NC1 was distinguished by enriched expression of *LGI4*, which is required for glia development and axon myelination ^69^. NC2 was predominantly derived from atrial tissues and expressed *GRIA1*, encoding glutamate receptor 1 which has altered expression levels in ischemic heart disease ^70^. Additionally, *GINS3* is enriched in these cells, which is known to participate in the regulation of cardiac repolarization ^71,72^. NC3 and NC4 subpopulations shared a broadly similar transcriptional profile including *LGR5*, a G-protein-coupled receptor involved in *Wnt*-signaling that promotes CM differentiation ^73^ and demarcates a subset of CM in the outflow tract ^74^; this region often becomes arrhythmogenic in heart disease ^75^. These cells also expressed genes associated with coronary artery disease, *PPP2R2B* ^76^, *LSAMP ^77^* and *LPL* an endothelial enzyme involved transporting lipoproteins into the heart ^78^. Additional smaller subpopulations (NC5-10) shared marker genes with other cell types and required further analyses to define their identities.

### Cross-tissue comparison between heart and skeletal muscle

We collected intercostal skeletal muscle samples from five healthy individuals, including one donor with matched cardiac tissue, and profiled the transcriptome of 35,665 single cells and 39,597 single nuclei. Analogous to the approach in the heart, the combination of cell- and nuclei-based methods allowed us to capture and resolve all major cell lineages, including myocyte and non-myocyte compartments (**Supplementary Figure G1A**).

We found that skeletal muscle had a similar distribution of cell populations to the heart, with a clear overlap of FB, EC with arterial and venous differentiation, PC and one major subpopulation of SMC (**Supplementary figure G2**). Cell-cell interaction analysis using CellPhoneDB.org predicted different interactions among arterial ECs and skeletal or heart muscle cells (**Figure 5A**). While *NOTCH* interactions were shared by both, EC-PC/SMC interactions in the heart were predicted to involve *JAG1/2 NOTCH3* and *NOTCH1 JAG1* but not in skeletal muscle. Additionally, the *EFNA1-EPHA3* receptor-ligand complex pair was consistently present in the heart but not detected in skeletal muscle. These differences may reflect distinct microvasculatures in the two tissues due to different oxygen requirements.

**Figure 5:**
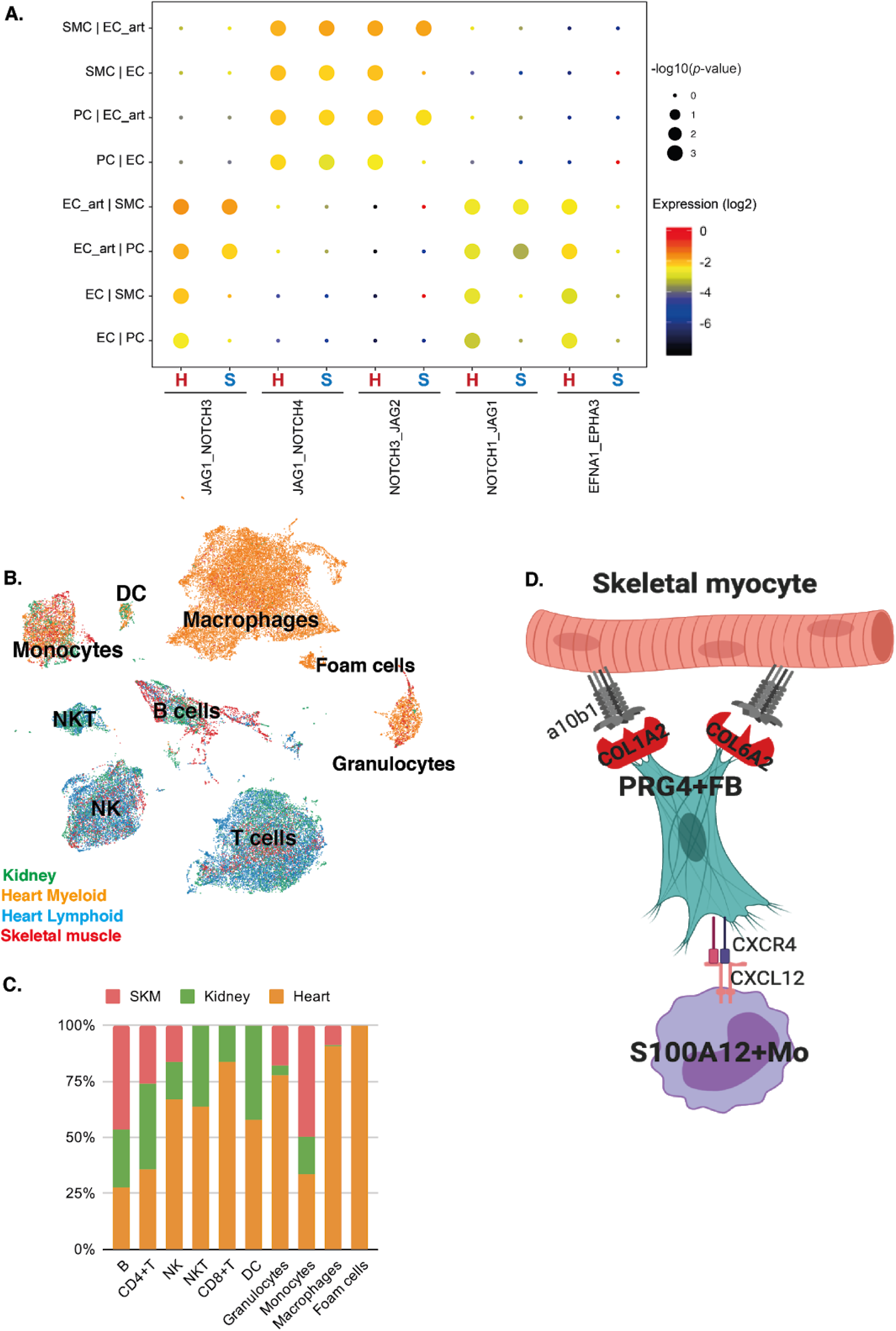
Cross-tissue comparison of cell populations and their interactions. **A**. Cell-cell interactions of vascular cells between skeletal muscle **(S)** and heart **(H)** showing that arterial EC from skeletal muscle share the same interactions as in the heart, while the skeletal muscle EC1 does not share the same interactions as their cardiac counterpart, and *EFNA1_EPHA3* is missing in the muscle. **B.** UMAP manifold comparing immune populations of the human heart, skeletal muscle and kidney, where cardiac macrophages are distinctive from the other tissues. **C**. Distribution of immune populations in the heart (myeloid and lymphoid), kidney and skeletal muscle. **D**. Schematic of the interaction between skeletal muscle and PRG4+FB mediated by type I collagen (COL1A2) and a10b1 integrins, and PRG4+ FB with S100A12+ monocytes mediated by CXCR4_CXCL12. The cell-cell interactions between the repair populations in the heart and skeletal muscle are very different. Created with BioRender ®.

We also compared the immune compartment of heart, skeletal muscle and adult kidney ^79^ by analyzing 40 206 cells and nuclei. Across the three tissues there was extensive overlap of lymphoid immune populations (B cells, CD4+ and CD8+ T cells, and NK/NKT cells). The overlap in myeloid populations (DC, granulocytes, foam cells, monocytes and macrophages) was not as extensive and several tissue resident macrophage populations had little overlap. We suggest that these tissue residents populations have developed transcriptional circuits tailored to the heart that differ from other tissues (**Figure 5B**, **5C** and **Supplementary Table G4**).

Myeloid cells in the skeletal muscle are known to interact with satellite cells to promote myogenesis ^80^. As we did not find a cardiac cell population analogous to skeletal muscle satellite cells in the heart, we considered if the heart had other potential repair mechanisms. To identify these we compared predicted interactions (**Supplementary Table G5**) among immune, fibroblast and myocyte populations in heart and skeletal muscle. These analyses showed that cardiac FB3 secrete FN1 and TNC which is bound by CM-expressed integrins. In skeletal muscle, predicted cell-cell interactions between *PRG4+*fibroblasts and skeletal muscle myocytes involved *COL1A2*, *COL6A2* and *a10b1* integrins (**Figure 5D**). Skeletal muscle fibroblasts and monocytes appear to interact *via* the *CXCR4_CXCL12* chemokine, while cardiomyocytes have a distinct interaction with macrophages as described above in **Figure 4D**. Altogether, these results imply interactive mechanisms are driven by different transcriptional circuits in heart and skeletal muscle. The circuit between CXCR4_CXCL12, which has been described to promote repair after myocardial infarction ^81^ appears to be primed by myeloid populations that initiate fibrotic repair.

### Association of GWAS hits with cardiac cell types

Using summary statistics of 12 well-powered GWAS studies of cardiac traits (**Supplementary Table H**) we applied the GWAS gene set enrichment method MAGMA^82,83^ to identify the most relevant cardiac cells for each trait, while removing redundant associations in conditional analyses. We found that atrial fibrillation (AF) GWAS signals were associated with gene expression profiles in vCM3, largely due to higher mean expression of *CAV1, CAV2, PRRX1* and other AF-associated genes in vCM3 cells. PR interval GWAS signals were associated with aCM population aCM5, with high expression *SCN5A, CAV1*, *ARHGAP24*, *MEIS1*, *TBX5* and *TTN*. GWAS signals for the QRS duration were associated with specific gene expression in NC1 (*PRKCA, CEP85L, SLC35F1, SIPA1L1, KLF12, FADS2*). Coronary artery disease (CAD) GWAS signals were associated with gene expression in a wide range of cell types (in particular SMC, FB, and EC). Similar results were also found for hypertension, in line with the disease relevance of vascular cells for both traits (**Supplementary Figure H**).

## Discussion

Our analyses of about half a million single cardiac cells and nuclei in six distinct cardiac regions from 14 donors significantly expand the characterisation of the heart’s cellular landscape, and contribute to a reference adult human heart cell atlas. By combining scRNA-Seq and snRNA-Seq data with state-of-the-art machine learning techniques, we provide detailed insights across the repertoire of cardiac cells, including CM (excluded by scRNA-Seq) as well as EC (underrepresented in cardiac snRNA-Seq) (**Figure 1C**). We quantify the cellular composition between cardiac chambers and across donors, highlighting chamber-specific features and differences between male and female donors.

Within each cell compartment we identified prototypic lineage-specific genes, differentially-expressed genes that characterized subpopulations with enrichment in particular chambers, and genes with previously unknown cardiac expression. We validated these findings using *in situ* smFISH with RNAscope probes (**Figure 2G, 3B; Supplementary Figure B1A**). Our results begin to unravel the molecular underpinnings of cardiac physiology and the multitudinous potential responses to stress and disease.

CM were the most prevalent cardiac cells and comprised higher percentages in ventricles than atria, as well as in left ventricular tissues from women *versus* men. As expected, aCM and vCM shared gene programs involved in excitation-contraction coupling, but also exhibit transcriptional differences indicative of the different developmental origins of these chambers from two primordial heart fields, and the vastly distinctive hemodynamic forces within adult atria and ventricles. We identified five different vCM subpopulations and six different aCM subpopulations, with variable proportions in the left and right chambers, and gene profiles of suggestive of canonical as well as specialized functions.

FB in atrial and ventricular tissues exhibited different transcriptional profiles and subpopulations, suggesting distinct functions. We found that ECM-producing and -organising fibroblasts, while present in atria and ventricles, differed in their mode of action, as exemplified by regional-specific expression of different collagen and ECM remodelling factor genes. Together with differences in the frequency of other cell types, we suggest that region-specific FB heterogeneity is critical to support CM across varying biophysical stimuli.

Cardiac EC are often under-studied due to their protruding nuclei, which may lead to damage during isolation. Indeed, in this study EC nuclei constituted 10.8% (atria) and 7.2% (ventricle) of cells, while immunohistochemistry data estimate that EC comprises >60% of non-myocytes in the heart ^84^. By sequencing single cells and single nuclei from the same sample, we resolved seven populations including arterial, venous and lymphatic EC, and an atrial-enriched population. Using CellPhoneDB.org ^85^, we inferred different arterial and venous EC interactions with mural cells.

A further cell compartment captured and analysed in depth were CD45+ immune cells. Due to their low frequency in tissues, the immune compartment is difficult to resolve using only nuclei. By performing CD45+ enrichment from the same cellular suspensions used for non-myocyte scRNA-Seq and using variational autoencoders, we identified 17 different immune cell populations across the myeloid and lymphoid lineages (**Figure 4A** and **4B**). This revealed novel cardiac-resident NK and NKT cells, which we show have analogous populations in human skeletal muscle and kidney. Specific to the heart are several interesting novel macrophage populations, including two anti-inflammatory (MØM2_tr*FOLR2*+ and MØ_tr*TMSB4X+*) macrophage populations which are predicted to be involved in heart repair (**Figure 4D**).

Assembling a reference cardiac cell atlas is of vital importance to understanding what goes awry in disease. The current coronavirus disease 2019 (COVID-19) pandemic, due to SARS-CoV-2, caused critical heart manifestations, ischemic symptoms with ST-elevation, reduced left ventricular contraction, and increased biomarkers for injury (NT-proBNP and troponins) ^86^. Related corona viral infections (SARS and MERS) also had cardiac involvement. Here, we found that expression of the viral receptor *ACE2* is higher in pericytes than in CM, as previously reported ^87^, but also that neither pericytes nor CM express the protease that prime viral entry, *TMPRSS2*. Instead CM and pericytes express *CTSB* and *CTSL*, which may also promote viral entry. *ACE2* expression correlated with *AGTR1* (angiotensin receptor-1) which was highest in pericytes, and consistent with its role of renin–angiotensin–aldosterone system (RAAS) signalling in cardiac hemodynamics. ACE2 cleaves the vasoconstrictor angiotensin II that binds AGTR. *ACE2*-null mice have reduced cardiac contractility, myocardial ischemia and hypoxia ^88, 89^ suggesting a profound role of ACE2 in the regulation of cardiovascular hemodynamics.

We expect that our results will allow further insights into other cardiac disease processes. Going forward, our results will furthermore be of value for deconvolution of existing bulk transcriptomic data, for transfer learning in analyses of other cardiac regions (valves, papillary muscle, conduction system), and to enable interpretation of the cellular responses to human heart disease. All of our data can be explored at www.heartcellatlas.org.

## Acknowledgments

This publication is part of the Human Cell Atlas - www.humancellatlas.org/publications. We gratefully acknowledge the Sanger Flow Cytometry Facility, Sanger Cellular Generation and Phenotyping (CGaP) Core Facility, and Sanger Core Sequencing pipeline for support with sample processing and sequencing library preparation. We thank J. Eliasova for graphical images, M. Prete and V. Kiselev for IT support, and S. Aldridge for editing the manuscript, and C. Domínguez-Conde, J. Park, K. James, K. Tuong and R. Elmentaite for helpful discussions about immune cell annotation, M. Luecken, M. Lotfollahi, D. Fischer and F. Theis for helpful discussions on computational analyses. M.N is the recipient of a British Heart Foundation (BHF) grant (PG/16/47/32156). M.N, N.H and S.A.T are funded by a BHF/DZHK grant (SP/19/1/34461). N.H is the recipient of an ERC Advanced Grant under the European Union Horizon 2020 Research and Innovation Program (grant agreement AdG788970) and the Federal Ministry of Education and Research of Germany in the framework of CaRNAtion (031L0075A). N.H, C.E.S, J.G.S are supported by grants from the Leducq Fondation (16CVD03). D.R received funding provided by the German Research Foundation (DFG). H.Z. is supported by the long-term Chinese Council Scholarship (CSC). M.K received funding provided by the Alexander von Humboldt Foundation. H.Z was supported by an EMBO long-term postdoctoral fellowship. G.O received funding provided by the CIHR Canadian Institutes for Health Research, HSF Heart and Stroke Foundation and AI Alberta Innovates. This project has been made possible in part by grant number 2019-202666 from the Chan Zuckerberg Initiative (C.E.S, J.G.S, M.N, N.H and S.A.T). C.T-L, K.P, E.F, E.T, K.R, O.B, H.Z, S.A.T are supported by the Wellcome Trust Sanger Institute grant (WT206194) and the Wellcome Science Strategic Support for a Pilot for the Human Cell Atlas (WT211276/Z/18/Z)). J.G.S and C.E.S are supported by NIH grants (2R01HL080494, 2R01HL080494, 1UM1HL098166). C.E.S and B.M are supported by the Howard Hughes Medical Institute. We are grateful to the deceased donors and their families and to the Cambridge Biorepository for Translational Medicine for access to human tissue.

## Data availability

Sequencing data will be made available upon publication via the Human Cell Atlas Data Coordination Platform. Data objects with the raw counts matrices and annotation will be made available upon publication via the www.heartcellatlas.org webportal.

## Author contributions

**Conceived the study**: N.H, S.A.T, M.N, C.E.S, J.G.S, H.M. **Acquired tissue**: K.M, K.S.P, G.O, H.Z, A.V **Processed tissue**: M.L, D.R, S.S, E.F, L.T, H.W, J.M.G, B.M, D.M.D, K.R, M.N. **Computational methods development:** C.T-L, K.P, M.H, J.G.S **Cell biology methods development:** M.L, S.S, E.L.L, M.N, H.M, D.R **Histopathological evaluation**: J.J.B, M.N **Data processing:** C.T-L, K.P, G.P **Global analysis:** C.T-L, D.R, M.L, C.L.W, E.L.L, J.G.S. **Cardiomyocyte analysis**: D.R, C.L.W, H.M, N.H, J.G.S, C.E.S. **Vascular analysis**: M.L, M.N, M.K, S.A.T. **Fibroblast analysis**: E.L.L, H.M, D.R, M.L, M.K, M.N, J.G.S, C.E.S, N.H. **Immune analysis**: C.T-L, S.A.T. **Neuronal analysis**: E.N, D.R, J.G.S, C.E.S. **Comparative analysis**: C.T-L, M.L, M.N, S.A.T. **GWAS analysis**: M.H, C.T-L, N.H, S.A.T. **Interpreted results**: M.L, C.T-L, D.R, H.M, C.L.W, E.L.L, M.H, J.B, M.N, C.E.S, J.G.S, S.A.T, N.H. **Wrote manuscript**: M.L, C.T-L, H.M, D.R, C.L.W, E.L.L, M.K, M.H, J.G.S, C.E.S, M.N, N.H, S.A.T.

## References

1. Fuchs, A. et al. Normal values of left ventricular mass and cardiac chamber volumes assessed by 320-detector computed tomography angiography in the Copenhagen General Population Study. Eur. Heart J. Cardiovasc. Imaging17, 1009–1017 (2016).

2. Meilhac, S. M. & Buckingham, M. E. The deployment of cell lineages that form the mammalian heart. Nat. Rev. Cardiol.15, 705–724 (2018).

3. Zhou, P. & Pu, W. T. Recounting Cardiac Cellular Composition. Circulation research vol. 118 368–370 (2016).

4. Massaia, A. et al. Single Cell Gene Expression to Understand the Dynamic Architecture of the Heart. Front Cardiovasc Med5, 167 (2018).

5. Lotfollahi, M., Wolf, F. A. & Theis, F. J. scGen predicts single-cell perturbation responses. Nat. Methods16, 715–721 (2019).

6. Chung Anne K. et al. Women Have Higher Left Ventricular Ejection Fractions Than Men Independent of Differences in Left Ventricular Volume. Circulation113, 1597–1604 (2006).

7. Ng, S. Y., Wong, C. K. & Tsang, S. Y. Differential gene expressions in atrial and ventricular myocytes: insights into the road of applying embryonic stem cell-derived cardiomyocytes for future therapies. Am. J. Physiol. Cell Physiol.299, C1234–49 (2010).

8. Johnson, E. K., Matkovich, S. J. & Nerbonne, J. M. Regional Differences in mRNA and lncRNA Expression Profiles in Non-Failing Human Atria and Ventricles. Sci. Rep.8, 13919 (2018).

9. Lee, J. H., Protze, S. I., Laksman, Z., Backx, P. H. & Keller, G. M. Human Pluripotent Stem Cell-Derived Atrial and Ventricular Cardiomyocytes Develop from Distinct Mesoderm Populations. Cell Stem Cell21, 179–194.e4 (2017).

10. Mazzotta, S. et al. Distinctive Roles of Canonical and Noncanonical Wnt Signaling in Human Embryonic Cardiomyocyte Development. Stem Cell Reports7, 764–776 (2016).

11. Shenton, F. C. & Pyner, S. Expression of transient receptor potential channels TRPC1 and TRPV4 in venoatrial endocardium of the rat heart. Neuroscience267, 195–204 (2014).

12. Delplancq, G. et al. Cardiomyopathy due to PRDM16 mutation: First description of a fetal presentation, with possible modifier genes. Am. J. Med. Genet. C Semin. Med. Genet. (2020) doi:10.1002/ajmg.c.31766.

13. Kim, S.-Y., Yasuda, S., Tanaka, H., Yamagata, K. & Kim, H. Non-clustered protocadherin. Cell Adh. Migr.5, 97–105 (2011).

14. Donlin, L. T. et al. Smyd2 controls cytoplasmic lysine methylation of Hsp90 and myofilament organization. Genes Dev.26, 114–119 (2012).

15. Gao, M. et al. Conserved expression of the PRELI domain containing 2 gene (Prelid2) during mid-later-gestation mouse embryogenesis. J. Mol. Histol.40, 227–233 (2009).

16. Denzel, M. S. et al. T-cadherin is critical for adiponectin-mediated cardioprotection in mice. J. Clin. Invest.120, 4342–4352 (2010).

17. Putku, M. et al. CDH13 promoter SNPs with pleiotropic effect on cardiometabolic parameters represent methylation QTLs. Hum. Genet.134, 291–303 (2015).

18. Liu, X. et al. Single-Cell RNA-Seq of the Developing Cardiac Outflow Tract Reveals Convergent Development of the Vascular Smooth Muscle Cells. Cell Rep.28, 1346–1361.e4 (2019).

19. Liu, R. & Jin, J.-P. Calponin isoforms CNN1, CNN2 and CNN3: Regulators for actin cytoskeleton functions in smooth muscle and non-muscle cells. Gene585, 143–153 (2016).

20. Besnard, S. et al. Smooth muscle dysfunction in resistance arteries of the staggerer mouse, a mutant of the nuclear receptor RORalpha. Circ. Res.90, 820–825 (2002).

21. Mikhailov, A. T. & Torrado, M. The enigmatic role of the ankyrin repeat domain 1 gene in heart development and disease. Int. J. Dev. Biol.52, 811–821 (2008).

22. Kwapiszewska, G. et al. Fhl-1, a new key protein in pulmonary hypertension. Circulation118, 1183–1194 (2008).

23. Geng, T. et al. H19 lncRNA Promotes Skeletal Muscle Insulin Sensitivity in Part by Targeting AMPK. Diabetes67, 2183–2198 (2018).

24. Huang, L. et al. Critical Roles of Xirp Proteins in Cardiac Conduction and Their Rare Variants Identified in Sudden Unexplained Nocturnal Death Syndrome and Brugada Syndrome in Chinese Han Population. J. Am. Heart Assoc.7, (2018).

25. Fittipaldi, S. et al. Alpha B-crystallin induction in skeletal muscle cells under redox imbalance is mediated by a JNK-dependent regulatory mechanism. Free Radic. Biol. Med.86, 331–342 (2015).

26. de Soysa, T. Y. et al. Single-cell analysis of cardiogenesis reveals basis for organ-level developmental defects. Nature572, 120–124 (2019).

27. Rajakumari, S. et al. EBF2 determines and maintains brown adipocyte identity. Cell Metab.17, 562–574 (2013).

28. 2016 AHA Late-Breaking Basic Science Abstracts. Circ. Res.119, e160–e171 (2016).

29. Bolotta, A. et al. New Insights into the Hepcidin-Ferroportin Axis and Iron Homeostasis in iPSC-Derived Cardiomyocytes from Friedreich’s Ataxia Patient. Oxid. Med. Cell. Longev.2019, 7623023 (2019).

30. Lakhal-Littleton, S. et al. An essential cell-autonomous role for hepcidin in cardiac iron homeostasis. Elife5, (2016).

31. Medioni, C. et al. Expression of Slit and Robo genes in the developing mouse heart. Dev. Dyn.239, 3303–3311 (2010).

32. Liu, C., Luo, N., Tung, C.-Y., Perrin, B. J. & Zhao, B. GRXCR2 Regulates Taperin Localization Critical for Stereocilia Morphology and Hearing. Cell Rep.25, 1268–1280.e4 (2018).

33. Wang, L. et al. Single-cell reconstruction of the adult human heart during heart failure and recovery reveals the cellular landscape underlying cardiac function. Nat. Cell Biol.22, 108–119 (2020).

34. Wei, Z. et al. Liprin-mediated large signaling complex organization revealed by the liprin-α/CASK and liprin-α/liprin-β complex structures. Mol. Cell43, 586–598 (2011).

35. Hoffmann, M. et al. SARS-CoV-2 Cell Entry Depends on ACE2 and TMPRSS2 and Is Blocked by a Clinically Proven Protease Inhibitor. Cell (2020) doi:10.1016/j.cell.2020.02.052.

36. Kalucka, J. et al. Single-Cell Transcriptome Atlas of Murine Endothelial Cells. Cell180, 764–779.e20 (2020).

37. Pober, J. S., Merola, J., Liu, R. & Manes, T. D. Antigen Presentation by Vascular Cells. Front. Immunol.8, 1907 (2017).

38. Corada, M., Morini, M. F. & Dejana, E. Signaling pathways in the specification of arteries and veins. Arterioscler. Thromb. Vasc. Biol.34, 2372–2377 (2014).

39. Zhao, Q. et al. Single-Cell Transcriptome Analyses Reveal Endothelial Cell Heterogeneity in Tumors and Changes following Antiangiogenic Treatment. Cancer Res.78, 2370–2382 (2018).

40. Ramachandran, P. et al. Resolving the fibrotic niche of human liver cirrhosis at single-cell level. Nature575, 512–518 (2019).

41. Kashiwazaki, M. et al. A high endothelial venule-expressing promiscuous chemokine receptor DARC can bind inflammatory, but not lymphoid, chemokines and is dispensable for lymphocyte homing under physiological conditions. Int. Immunol.15, 1219–1227 (2003).

42. Xiao, Y. et al. Hippo Signaling Plays an Essential Role in Cell State Transitions during Cardiac Fibroblast Development. Dev. Cell45, 153–169.e6 (2018).

43. Awwad, K. et al. Role of secreted modular calcium-binding protein 1 (SMOC1) in transforming growth factor β signalling and angiogenesis. Cardiovasc. Res.106, 284–294 (2015).

44. Tang, J. et al. Genetic Fate Mapping Defines the Vascular Potential of Endocardial Cells in the Adult Heart. Circ. Res.122, 984–993 (2018).

45. Barberis, M. C., Faleri, M., Veronese, S., Casadio, C. & Viale, G. Calretinin. A selective marker of normal and neoplastic mesothelial cells in serous effusions. Acta Cytol.41, 1757–1761 (1997).

46. Zhou, B. et al. Adult mouse epicardium modulates myocardial injury by secreting paracrine factors. J. Clin. Invest.121, 1894–1904 (2011).

47. Gambardella, L. et al. BNC1 regulates cell heterogeneity in human pluripotent stem cell-derived epicardium. Development146, (2019).

48. Stallcup, W. B. The NG2 Proteoglycan in Pericyte Biology. Adv. Exp. Med. Biol.1109, 5–19 (2018).

49. Vanlandewijck, M. et al. A molecular atlas of cell types and zonation in the brain vasculature. Nature554, 475–480 (2018).

50. Lee, J.-H. et al. Anatomically and Functionally Distinct Lung Mesenchymal Populations Marked by Lgr5 and Lgr6. Cell170, 1149–1163.e12 (2017).

51. Daniel, J.-M. et al. Regulator of G-Protein Signaling 5 Prevents Smooth Muscle Cell Proliferation and Attenuates Neointima Formation. Arterioscler. Thromb. Vasc. Biol.36, 317–327 (2016).

52. Wong, A. P., Nili, N. & Strauss, B. H. In vitro differences between venous and arterial-derived smooth muscle cells: potential modulatory role of decorin. Cardiovasc. Res.65, 702–710 (2005).

53. Sweeney, M. & Foldes, G. It Takes Two: Endothelial-Perivascular Cell Cross-Talk in Vascular Development and Disease. Front Cardiovasc Med5, 154 (2018).

54. Chen, Q. et al. Endothelial cells are progenitors of cardiac pericytes and vascular smooth muscle cells. Nat. Commun.7, 12422 (2016).

55. Ferland-McCollough, D., Slater, S., Richard, J., Reni, C. & Mangialardi, G. Pericytes, an overlooked player in vascular pathobiology. Pharmacol. Ther.171, 30–42 (2017).

56. Guimarães-Camboa, N. et al. Pericytes of Multiple Organs Do Not Behave as Mesenchymal Stem Cells In Vivo. Cell Stem Cell20, 345–359.e5 (2017).

57. Burstein Brett, Libby Eric, Calderone Angelino & Nattel Stanley. Differential Behaviors of Atrial Versus Ventricular Fibroblasts. Circulation117, 1630–1641 (2008).

58. Hanna, N., Cardin, S., Leung, T.-K. & Nattel, S. Differences in atrial versus ventricular remodeling in dogs with ventricular tachypacing-induced congestive heart failure. Cardiovasc. Res.63, 236–244 (2004).

59. Zhang, F., Li, C., Halfter, H. & Liu, J. Delineating an oncostatin M-activated STAT3 signaling pathway that coordinates the expression of genes involved in cell cycle regulation and extracellular matrix deposition of MCF-7 cells. Oncogene22, 894–905 (2003).

60. Ghatak, S. et al. Transforming growth factor β1 (TGFβ1) regulates CD44V6 expression and activity through extracellular signal-regulated kinase (ERK)-induced EGR1 in pulmonary fibrogenic fibroblasts. J. Biol. Chem.292, 10465–10489 (2017).

61. Abe, H. et al. Macrophage hypoxia signaling regulates cardiac fibrosis via Oncostatin M. Nat. Commun.10, 2824 (2019).

62. Dick, S. A. et al. Self-renewing resident cardiac macrophages limit adverse remodeling following myocardial infarction. Nat. Immunol.20, 29–39 (2019).

63. Yoo, W. et al. Progranulin attenuates liver fibrosis by downregulating the inflammatory response. Cell Death Dis.10, 758 (2019).

64. Adamo, L., Rocha-Resende, C., Prabhu, S. D. & Mann, D. L. Reappraising the role of inflammation in heart failure. Nat. Rev. Cardiol. (2020) doi:10.1038/s41569-019-0315-x.

65. Kim, D. H. et al. Novel Role of Lck in Leptin-Induced Inflammation and Implications for Renal Aging. Aging Dis.10, 1174–1186 (2019).

66. Efremova, M., Vento-Tormo, M., Teichmann, S. A. & Vento-Tormo, R. CellPhoneDB: inferring cell-cell communication from combined expression of multi-subunit ligand-receptor complexes. Nat. Protoc. (2020) doi:10.1038/s41596-020-0292-x.

67. Herring, N., Kalla, M. & Paterson, D. J. The autonomic nervous system and cardiac arrhythmias: current concepts and emerging therapies. Nat. Rev. Cardiol.16, 707–726 (2019).

68. Bhattacharyya, S. & Munshi, N. V. Development of the Cardiac Conduction System. Cold Spring Harb. Perspect. Biol. (2020) doi:10.1101/cshperspect.a037408.

69. Kegel, L., Aunin, E., Meijer, D. & Bermingham, J. R. LGI proteins in the nervous system. ASN Neuro5, 167–181 (2013).

70. Gronich, N., Kumar, A., Zhang, Y., Efimov, I. R. & Soldatov, N. M. Molecular remodeling of ion channels, exchangers and pumps in atrial and ventricular myocytes in ischemic cardiomyopathy. Channels4, 101–107 (2010).

71. Newton-Cheh, C. et al. Common variants at ten loci influence QT interval duration in the QTGEN Study. Nat. Genet.41, 399–406 (2009).

72. Milan, D. J. et al. Drug-sensitized zebrafish screen identifies multiple genes, including GINS3, as regulators of myocardial repolarization. Circulation120, 553–559 (2009).

73. Jha, R. et al. Downregulation of LGR5 Expression Inhibits Cardiomyocyte Differentiation and Potentiates Endothelial Differentiation from Human Pluripotent Stem Cells. Stem Cell Reports9, 513–527 (2017).

74. Sahara, M. et al. Population and Single-Cell Analysis of Human Cardiogenesis Reveals Unique LGR5 Ventricular Progenitors in Embryonic Outflow Tract. Dev. Cell48, 475–490.e7 (2019).

75. Lerman, B. B. Outflow tract ventricular arrhythmias: An update. Trends Cardiovasc. Med.25, 550–558 (2015).

76. Nolan, D. K. et al. Fine mapping of a linkage peak with integration of lipid traits identifies novel coronary artery disease genes on chromosome 5. BMC Genet.13, 12 (2012).

77. Wang, L. et al. Polymorphisms of the tumor suppressor gene LSAMP are associated with left main coronary artery disease. Ann. Hum. Genet.72, 443–453 (2008).

78. Olivecrona, G. Role of lipoprotein lipase in lipid metabolism. Curr. Opin. Lipidol.27, 233–241 (2016).

79. Stewart, B. J. et al. Spatiotemporal immune zonation of the human kidney. Science365, 1461–1466 (2019).

80. Madaro, L. et al. Macrophages fine tune satellite cell fate in dystrophic skeletal muscle of mdx mice. PLoS Genet.15, e1008408 (2019).

81. Döring, Y., Pawig, L., Weber, C. & Noels, H. The CXCL12/CXCR4 chemokine ligand/receptor axis in cardiovascular disease. Front. Physiol.5, 212 (2014).

82. de Leeuw, C. A., Mooij, J. M., Heskes, T. & Posthuma, D. MAGMA: generalized gene-set analysis of GWAS data. PLoS Comput. Biol.11, e1004219 (2015).

83. Watanabe, K., Umićević Mirkov, M., de Leeuw, C. A., van den Heuvel, M. P. & Posthuma, D. Genetic mapping of cell type specificity for complex traits. Nat. Commun.10, 3222 (2019).

84. Pinto, A. R. et al. Revisiting Cardiac Cellular Composition. Circ. Res.118, 400–409 (2016).

85. Vento-Tormo, R. et al. Single-cell reconstruction of the early maternal-fetal interface in humans. Nature563, 347–353 (2018).

86. Huang, C. et al. Clinical features of patients infected with 2019 novel coronavirus in Wuhan, China. Lancet395, 497–506 (2020).

87. Chen, L., Li, X., Chen, M., Feng, Y. & Xiong, C. The ACE2 expression in human heart indicates new potential mechanism of heart injury among patients infected with SARS-CoV-2. Cardiovasc. Res. (2020) doi:10.1093/cvr/cvaa078.

88. Krege, J. H. et al. Male-female differences in fertility and blood pressure in ACE-deficient mice. Nature375, 146–148 (1995).

89. Crackower, M. A. et al. Angiotensin-converting enzyme 2 is an essential regulator of heart function. Nature417, 822–828 (2002).

