## Supplementary Material for "Cells and gene expression programs in the adult human heart" (from another version of this preprint: version 1, 2020-04-05)

**Experimental methods:**

*Research ethics for donor tissues:* Heart tissues (D1-7 and 11) processed at Wellcome Sanger Institute (Hinxton, UK) were obtained from deceased transplant organ donors after Research Ethics Committee approval (Ref 15/EE/0152, East of England - Cambridge South Research Ethics Committee) and informed consent from the donor families. Heart tissues (H2-7) processed at Harvard Medical School (Boston, MA, USA) were obtained from deceased organ donors after Human Research Ethics Board approval Pro00011739 (University of Alberta, Edmonton, Canada). Informed consent from donor families was acquired *via* the institutional Human Organ Procurement and Exchange Program (HOPE). Cardiovascular history was unremarkable for all donors (**Supplementary Table A1**).

*Tissue acquisition and processing:* For Cambridge/UK donors (D1-7 and 11), full-thickness myocardial biopsies from LA, RA, RV, SP, AX and LV were collected from donor hearts (**Supplementary Table A1**), and stored in ice-cold cardioplegic solution. Samples were transported and kept on ice at all times. Single cell isolation and CD45+ enrichment were carried out on freshly collected samples. The remaining tissue was formalin-fixed or flash-frozen as indicated below and stored at -80°C. For North American donors (H2-7), cold cardioplegia through the aortic root was employed before heart excision, followed by storage in ice-cold saline solution during transportation to the laboratory. Full-thickness myocardial biopsies from LA, RA, RV, SP, AX and LV were utilized by avoiding epicardial fat. All samples were flash-frozen and stored at -80°C.

*Single nuclei isolation:* Single nuclei were obtained from flash-frozen tissues using mechanical homogenization as previously described [1]. Tissues were homogenized using a 7ml glass Dounce tissue grinder set (Merck) with 8-10 strokes of a loose pestle (A) and 8-10 strokes of a tight pestle (B) in homogenization buffer (250mM Sucrose, 25mM KCl, 5mM MgCl2, 10mM Tris-HCl, 1mM dithiothreitol (DTT), 1x Protease Inhibitor, 0.4U/μl RNaseIn, 0.2U/μl SUPERaseIn, 0.1% Triton X-100 in Nuclease-Free Water). Homogenate was filtered through a 40μm cell strainer (Corning). After centrifugation (500xg, 5 min, 4°C) the supernatant was removed and the pellet was resuspended in storage buffer (1x Phosphate-Buffered Saline (PBS), 4% Bovine Serum Albumin (BSA), 0.2U/μl Protector RNaseIn). Nuclei were stained with NucBlue Live ReadyProbes Reagents (ThermoFisher) and Hoechst-positive single nuclei were purified by fluorescent activated cell sorting using

Influx, XDP or FACSAria (BD Biosciences). Nuclei purification and integrity was verified under microscope and they were further processed using the Chromium controller (10X Genomics) according to the manufacturer's protocol.

*Single cell preparation:* Heart tissues (0.2-0.9g) were transferred from cardioplegic solution into gentleMACS C-tubes (Miltenyi Biotec) containing enzymatic digestion base solution (HBSS, 10mM HEPES, 30mM Taurin). Tissues were minced using scissors (FST) and automatically digested using gentleMACS Octo Dissociator (Miltenyi Biotec) with heaters. Cardiomyocyte depleted single-cell suspension was washed with base solutioncontaining 20% Fetal Bovine Serum (FBS, Gibco), filtered through 70µm nylon strainer (BD Falcon), collected by centrifugation (330g, 10min, 4°C) and resuspended in base solution containing 0.2% FBS (Gibco). Cells were manually counted three times by Trypan blue exclusion after each centrifugation and resuspended at a concentration of ≥ $2x10^6$/ml [2]. Single cells were processed using Chromium controller (10X Genomics) according to the manufacturer's protocol.

*CD45+ cell enrichment:* Cell suspension was prepared as described above. For CD45+ enrichment cells were labelled using anti-human CD45 monoclonal antibody-conjugated microbeads according to the manufacturer’s protocol (Miltenyi Biotec). Briefly, up to $10^7$ cells were incubated for 15 min at 4°C in 80µl of buffer (1x PBS pH 7.2, 0.5% BSA, 2mM EDTA) containing 20µl CD45 microbeads. Cell suspension was washed in PBS/BSA/EDTA buffer once and pelleted by centrifugation (330g, 10min, 4°C). Resuspended cells were applied to MACS LS columns (Miltenyi Biotec). CD45-depleted cell fraction was discarded after three washes with PBS/BSA/EDTA buffer and CD45+ cell fraction was collected in PBS/BSA/EDTA buffer by removal of the columns from the magnetic field. CD45+ cells were counted and resuspended in PBS/BSA/EDTA buffer to a concentration of ≥ $2x10^6$/ml before further processing using a Chromium controller (10X Genomics) according to the manufacturer's protocol.

*Chromium 10X library preparation:* Single cells and nuclei were manually counted by Trypan blue exclusion or automatically using a Countess II (Life technologies) using at least two separate counts. Cell or nuclei suspension was adjusted to 400-1,000 cells/µl and loaded on the Chromium controller (10X Genomics) with a targeted cell/nuclei recovery of 4,000-10,000 per run. 3’ gene expression libraries were prepared according to the manufacturer's instructions of the v2 or v3 Chromium Single Cell Reagent Kits (10X Genomics). Quality control of cDNA and final libraries were done using Bioanalyzer High

Sensitivity DNA Analysis (Agilent) or 4200 TapeStation System (Agilent). Libraries were sequenced using HiSeq4000 (Illumina) at Wellcome Sanger Institute, and Nextseq500 (Illumina) at Harvard Medical School with a minimum depth of 20,000-30,000 read pairs per cell or nucleus.

*Spatial validation using smFISH with RNAscope probes:* During preparation of formalin-fixed paraffin-embedded (FFPE) samples fresh tissue was fixed in neutral-buffered 10% formalin for 18-36 hrs and subsequently embedded in paraffin blocks. Fixed-frozen (FF) tissue samples were fixed in 4% paraformaldehyde (ThermoFisher). Sections were cut at 5µm thickness using a microtome, and placed onto SuperFrost Plus slides (VWR). FFPE tissue slides were automatically stained using BOND RX (Leica) and the RNAscope Multiplex Fluorescent Reagent Kit v2 Assay (ACDBio) according to the manufacturer's protocol. FF tissue slides were processed according to the protocol of RNAscope Multiplex Fluorescent Assay v1 (ACDBio). RNAscope ready- or custom-made target probes were run in parallel to multiplex positive and negative controls (**Supplementary Table I3**). All nuclei were DAPI-stained. FFPE tissue slides were imaged using an Opera Phenix High-Content confocal Screening System (Perkin Elmer) and 20x water-immersion objective. FF tissue slides were imaged using a LSM710 confocal microscope (Zeiss) and 40x oil objective. Visualization and background removal (rolling ball radius) was done using Fiji/ImageJ [3].

*Hematoxylin&Eosin staining:* Tissue samples were fresh-frozen in isopentane (ThermoFisher) at -80°C and OCT (VWR) embedded. Sections were cut at a thickness of 10µm using a microtome, placed onto SuperFrostPlus slides (VWR) and further processed according to standard Hematoxylin&Eosin staining protocol (**Supplementary Figure A1**).

*Skeletal muscle tissue processing*

*Tissue acquisition:* Intercostal muscle samples were obtained from between the 2nd and 3rd rib on the left side. This is typically from the deepest layer of muscle (furthest away from the skin). Samples were collected directly into cold preservation solution.

*Nuclei isolation:* Muscle tissue was washed in 1x PBS, cleaned of any visible fat depositions and minced to obtain fragments of approximately $1mm^3$. Per each sample, ~0.3g of minced tissues was homogenised in 3ml of Buffer A (250mM sucrose, 10mg/ml BSA, 5mM $MgCl_2$, 0.12U/µl RNAseIn, 0.06U/µl SUPERasIn, 1x Protease Inhibitor) using Dounce tissue grinder set (Merck) with 50 strokes of the loose pestle (A). The homogenate was filtered through a 100µm cell strainer (Corning) and the strainer was washed twice with 1ml and with 750µl of

Buffer A. After adding Triton X-100 (final concentration 0.5%), the mixture was further homogenised with 50 strokes of the tight pestle (B). Following filtering through a 40μm strainer, nuclei were centrifuged (3000g, 5min, 4°C), resuspended in 1ml of Buffer B (320mM Sucrose, 10mg/ml BSA, 3mM CaCl2, 2mM MgAc2, 0.1mM EDTA, 10mM Tris-HCl, 1mM DTT, 1x Protease Inhibitor, 0.12U/μl RNAseIn, 0.06U/μl SUPERasin) and purified using a 27% percoll gradient solution. The percoll mixture was centrifuged at (20,000g, 15min, 4℃) and the pellet resuspended in 200μL of Buffer B, followed by centrifugation (20,000g, 3min, 4℃). After Trypan Blue staining, the intact nuclei were counted using a hemocytometer. Nuclei were profiled using a Chromium controller (10X Genomics) according to the manufacturer's protocol.

*Single cell isolation*: Muscle tissue was washed in 1X PBS, cleaned of any visible fat depositions and finely minced. 2g of the minced tissue was transferred to digestion buffer 1 (750U/ml Collagenase type 2 in 1x PBS) and incubated at 37℃ in a water bath for 90 min. The partially digested tissue was collected by centrifugation (650g, 5min, 4℃) and the pellet resuspended in digestion buffer 2 (100U/ml Collagenase type 2, 2U/ml Dispase in PBS). After 30 min incubation at 37℃ in a water bath, the digestion was stopped by adding 2% FBS. Cells were filtered through a 100μm and a 40μm nylon strainer (BD Falcon), collected by centrifugation (650g, 4℃, 3 min) and washed with 1x PBS/ 2% FBS. Subsequently, 20% percoll gradient (15,000g, 4 ℃, 20 min) was used for cells' purification. The layer containing cells was collected, washed in PBS containing 2% FBS and viable cells were counted by Trypan Blue exclusion using a hemocytometer. Nuclei were profiled using a Chromium controller (10X Genomics) according to the manufacturer's protocol.

The methods key resources table is found in **Supplementary Table I1**.

**Data analysis:**

*Transcriptome mapping:* After sequencing, samples were demultiplexed and stored as CRAM files. Each sample was mapped to the human reference genome (GRCh38 v3.0.0) provided by 10X Genomics, and using the CellRanger suite (v3.0.1) with default parameters. Single cell samples were mapped against the reference as it was provided. Single nuclei samples, the reference for *pre-mRNA* was created using the 10X Genomics instructions (https://support.10xgenomics.com/single-cell-gene-expression/software/pipelines/latest/advanced/references).

*Count data processing*: After mapping, samples from each data source (single nuclei, single cell and CD45+ cell) were grouped into individual AnnData objects by concatenating the raw_feature_bc_matrix_h5.h5 and adding the appropriate metadata information. For each data source object, the mean of UMIs (n_counts) was calculated and used as a threshold for empty droplets.

*Doublet detection*: After empty droplets removal, we applied scrublet[4] to assign a doublet score (scrublet_score) to each cell. These cells were clustered and visualised using the UMAP method [5]. Additionally, we processed each cell for doublet detection using a percolation method to allow for improved detection of doublets [6].

*Background removal:* Background contamination removal was performed using CellBender [7] using 25 000 empty barcodes, 300 epochs and the number of recovered cells for each single cell run.

*Cell QC and filtering*: Each data source was processed and annotated separately to account for source-specific quality differences. These metrics were included as covariates for further processing. Total cells and CD45+ cells were filtered for counts (500 < n_counts < 15000), genes (200 < n_genes), mitochondrial genes (percent_mito < 20%), ribosomal genes (percent_ribo < 20%) and scrublet score (scrublet_score < 0.3). Single nuclei were filtered for counts (500 < n_counts < 15000), genes (300 < n_genes < 6000), mitochondrial genes (percent_mito < 5%), ribosomal genes (percent_ribo < 5%) and scrublet score (scrublet_score < 0.3).

*Scanpy* toolkit 1.4 [8] in Python version 3.7 was used to perform downstream analyses, including normalization (normalize_per_cell: counts_per_cell_after = 10,000), log transformation (log1p), variable gene detection (highly_variable_genes), regressing out unwanted sources of variation (regress_out: n_counts and percent_mito), data feature scaling (scale: max_value = 10) and PCA (pca: using highly variable genes) as previously described [9].

*Batch alignment using deep variational autoencoders*: We built a global manifold by aligning all the data sources and donors in our data. This was done in a two-step procedure:

a) Each source is analysed and annotated separately aligning only for donors using a PC-space linear regression step prior to batch alignment with *bbknn* [10]. DEGs were calculated using a Wilcoxon Rank Sum Test with Bonferroni-Hochberg adjustment as implemented in the *Scanpy* framework.

b) To annotate each cluster, we used an integrative approach by searching the top significant DEGs (*p*-value < 1*e*-05) with a logFC > 1 against the *ToppFun* [11] and *EnrichR* [12] databases. Significant hits on *Pathways*, *Transcriptional regulation* and *Biological processes* were prioritised to annotate a given cluster. Each cellular compartment was labelled under the adata.obs['cell_type'] slot after grouping source-specific cell states.
c) All sources were combined into a single AnnData object under the label adata.obs['cell_sources']. Batches were aligned using the *batch_correction* function from the *scGen* variational autoencoder [13]. First we aligned for adata.obs['cell_sources'], using adata.obs['cell_type'] as an anchor. Next we aligned for adata.obs['donor'], using adata.obs['cell_type'] as an anchor. Each batch alignment round was run for 50 epochs.

The final global manifold was stored in an AnnData object with raw counts in the main matrix and CellBender-processed counts in the raw matrix. Manifolds for the fibroblasts, vascular, immune cardiac populations, as well as the skeletal muscle analysis were created using this method.

*NOTE*: Initially, we used the CellBender-processed-matrix to calculate with DEG in order to avoid any background contamination to interfere with the cluster annotation. Nevertheless, the background removal was not satisfactory, so we used the cellranger count matrix instead.

*Cell-Cell interactions*: Expression matrices of the populations under study were exported from the AnnData, together with a metadata table that contain the cell-barcodes as indices. We then run CellPhoneDB as follows: cellphonedb method statistical_analysis meta.tsv counts.tsv --subsampling –subsampling-log True –subsampling-num-cells 50000 --counts-data=gene_name --threads = 30.

*Estimation of RNA Velocity*: To calculate the RNA velocity of the single cells and CD45+ enriched single cells, we used the CellRanger output BAM file and the GENCODE v33 GTF (ftp://ftp.ebi.ac.uk/pub/databases/gencode/Gencode_human/release_33/gencode.v33.chr_patch_hapl_scaff.annotation.gtf.gz) file together with the *velocyto*[14] CLI v0.17.17 to generate a loom file containing the quantification of spliced and unspliced RNA. Next, we built a manifold, clustered the cells and visualised the RNA velocities using *scVelo*[15].

*Subpopulation analyses of aCM, vCM, aFB, vFB and NC:* All nuclei labelled in the global object as cardiomyocytes, fibroblasts and neural cells were selected for further subpopulation analyses. Additional cell population-specific filtering criteria were applied as follows: CM & NC - counts (n_counts < 12500), genes (n_genes < 4000), mitochondrial genes (percent_mito < 1%), ribosomal genes (percent_ribo < 1%) and scrublet score (scrublet_score < 0.25); FB - counts (n_counts < 7500), genes (n_genes < 3000), mitochondrial genes (percent_mito < 1%), ribosomal genes (percent_ribo < 2%). For CM and FB the resulting nuclei were then further split into two groupings based on the region of origin: i) LA and RA and ii) LV, RV, AX and SP.

Donor effects were aligned as described in step a) above. For regional subclustering of FB and NC we used *harmony* for the harmonization of donor-specific gene expression patterns [16]. Leiden and Louvain (NC clustering) clustering and UMAP visualization were performed for identifying subpopulations and visualization[17]. Differentially expressed genes were calculated using the Wilcoxon Rank-Sum test. Genes were ranked by score.

*Gene Ontology enrichment analysis:* We used the R package gProfileR (https://cran.r-project.org/web/packages/gProfileR/index.html) with the score-ranked gene list as input and the set of genes expressed in vCM as background (those genes having a UMI count > 1).

*Gene set score*: We used the score_genes function as implemented in *scanpy* to calculate the enrichment of genes involved in the Oncostatin M pathway. A list of genes was collected upon literature research [18,19]. For gene set enrichment, only highly expressed genes were considered to reduce noise (more than 500 UMIs across all cells).

*Statistics:* All analyses were performed using *R Software, Version 3.6.1.* Student t-tests (t-tests) were used to compare cell type distributions at each site. A *p*-value<0.05 was considered statistically significant. Linear regression models (correlations) were obtained using the R linear model function (lm), which estimated statistical likelihood (p-value) of a linear relationship. Bonferroni correction was applied for multiple testing.

GWAS enrichment analysis:

We downloaded GWAS summary statistics from broad cvdi, EBI GWAS catalog and GWAS atlas. We selected traits with well powered GWAS (N > 5000 and number of significant loci > 10). GWAS data sets are summarized in (**Supplementary Table H1)**. Gene expression data of protein coding genes were mapped onto Entrez gene ids and these gene annotations were used on the human genome assembly hg19 / 37. We only used gene expression data from nuclei. We implemented the analysis described in [20] in python and R. The log transformed counts were used to compute average cell type specific expression profiles. We used the results from the subpopulation annotations from the subclustering analyses. We performed individual magma analyses for each cell type, always condition on default gene level covariates (e.g. gene length) and average gene expression across all cells. Subsequently we applied the Benjamini Hochberg method and selected cell type trait associations with FDR < 10%. These pairs were then subjected to conditional analysis as described in [20] to define "independent", "jointly explained" and "partially jointly explained" pairs of associations.

**References**:


1. Litvinukova, M. *et al.* Single Cell and Single Nuclei Analysis Human Heart Tissue v1 (protocols.io.veae3ae). doi:10.17504/protocols.io.veae3ae.
2. Noseda, M. *et al.* PDGFRα demarcates the cardiogenic clonogenic Sca1+ stem/progenitor cell in adult murine myocardium. *Nat. Commun.* **6**, 6930 (2015).
3. Schindelin, J. *et al.* Fiji: an open-source platform for biological-image analysis. *Nat. Methods* **9**, 676–682 (2012).
4. Wolock, S. L., Lopez, R. & Klein, A. M. Scrublet: Computational Identification of Cell Doublets in Single-Cell Transcriptomic Data. *Cell Syst* **8**, 281–291.e9 (2019).
5. McInnes, L. & Healy, J. UMAP: Uniform Manifold Approximation and Projection for Dimension Reduction. *arXiv [stat.ML]* (2018).
6. Popescu, D.-M. *et al.* Decoding human fetal liver haematopoiesis. *Nature* **574**, 365–371 (2019).

7. Fleming, S. J., Marioni, J. C. & Babadi, M. CellBender remove-background: a deep generative model for unsupervised removal of background noise from scRNA-seq datasets. *bioRxiv* 791699 (2019) doi:10.1101/791699.
8. Wolf, F. A., Angerer, P. & Theis, F. J. SCANPY: large-scale single-cell gene expression data analysis. *Genome Biol.* **19**, 15 (2018).
9. Luecken, M. D. & Theis, F. J. Current best practices in single-cell RNA-seq analysis: a tutorial. *Mol. Syst. Biol.* **15**, e8746 (2019).
10. Polański, K. *et al.* BBKNN: Fast Batch Alignment of Single Cell Transcriptomes. *Bioinformatics* (2019) doi:10.1093/bioinformatics/btz625.
11. Chen, J., Bardes, E. E., Aronow, B. J. & Jegga, A. G. ToppGene Suite for gene list enrichment analysis and candidate gene prioritization. *Nucleic Acids Res.* **37**, W305–11 (2009).
12. Kuleshov, M. V. *et al.* Enrichr: a comprehensive gene set enrichment analysis web server 2016 update. *Nucleic Acids Res.* **44**, W90–7 (2016).
13. Lotfollahi, M., Wolf, F. A. & Theis, F. J. scGen predicts single-cell perturbation responses. *Nat. Methods* **16**, 715–721 (2019).
14. La Manno, G. *et al.* RNA velocity of single cells. *Nature* **560**, 494–498 (2018).
15. Bergen, V., Lange, M., Peidli, S., Alexander Wolf, F. & Theis, F. J. Generalizing RNA velocity to transient cell states through dynamical modeling. *bioRxiv* 820936 (2019) doi:10.1101/820936.
16. Korsunsky, I. *et al.* Fast, sensitive and accurate integration of single-cell data with Harmony. *Nat. Methods* **16**, 1289–1296 (2019).
17. Traag, V. A., Waltman, L. & van Eck, N. J. From Louvain to Leiden: guaranteeing well-connected communities. *Sci. Rep.* **9**, 5233 (2019).
18. Dey, G. *et al.* Signaling network of Oncostatin M pathway. *J. Cell Commun. Signal.* **7**, 103–108 (2013).

19. Abe, H. *et al.* Macrophage hypoxia signaling regulates cardiac fibrosis via Oncostatin M. *Nat. Commun.* **10**, 2824 (2019).

20. Watanabe, K., Umićević Mirkov, M., de Leeuw, C. A., van den Heuvel, M. P. & Posthuma, D. Genetic mapping of cell type specificity for complex traits. *Nat. Commun.* **10**, 3222 (2019).
